# Identification of a Druggable Site on GRP78 at the GRP78-SARS-CoV-2 Interface and Compounds to Disrupt that Interface

**DOI:** 10.1101/2023.09.12.557363

**Authors:** Maria Lazou, Jonathan R. Hutton, Arijit Chakravarty, Diane Joseph-McCarthy

## Abstract

SARS-CoV-2, the virus that causes COVID-19, led to a global health emergency that claimed the lives of millions. Despite the widespread availability of vaccines, the virus continues to exist in the population in an endemic state which allows for the continued emergence of new variants. Most of the current vaccines target the spike glycoprotein interface of SARS-CoV-2, creating a selection pressure favoring viral immune evasion. Antivirals targeting other molecular interactions of SARS-CoV-2 can help slow viral evolution by providing orthogonal selection pressures on the virus. GRP78 is a host auxiliary factor that mediates binding of the SARS-CoV-2 spike protein to human cellular ACE2, the primary pathway of cell infection. As GRP78 forms a ternary complex with SARS-CoV-2 spike protein and ACE2, disrupting the formation of this complex is expected to hinder viral entry into host cells. Here, we developed a model of the GRP78-spike protein-ACE2 complex. We then used that model together with hot spot mapping of the GRP78 structure to identify the putative binding site for spike protein on GRP78. Next, we performed structure-based virtual screening of known drug/candidate drug libraries to identify binders to GRP78 that are expected to disrupt spike protein binding to the GRP78, and thereby preventing viral entry to the host cell. A subset of these compounds have previously been shown to have some activity against SARS-CoV-2. The identified hits are starting points for the further development of novel SARS-CoV-2 therapeutics, potentially serving as proof-of-concept for GRP78 as a potential drug target for other viruses.

## Introduction

To date, more than half of a billion people have been infected with COVID-19, a disease caused by the SARS-CoV-2 virus. As of December 2021, the disease was responsible for around 15 to 18 million excess deaths and has affected the lives of every human through government shutdowns and economic recessions[1]. The virus has evolved rapidly since its emergence in humans in late 2019, with untold numbers of variants emerging around the world[2]. Due to its incredibly high rate of transmission and propagation, SARS-CoV-2 variants evolve rapidly and continue to pose a serious threat to treatment effectiveness despite the availability of vaccines [2, 3]. The major pathway through which many coronaviruses, including SARS-CoV-2, infect the cell is by binding the spike protein to cell-surface angiotensin-converting enzyme 2 (ACE2), resulting in endocytosis of the complex into the cell[4, 5]. Antibody therapeutics, including vaccines, directly target this spike-ACE2 interface to block viral infection[6–8]. However, it has been shown that the SARS-CoV-2 spike protein has a large amount of evolutionary space—mutations that affect the binding of neutralizing antibodies to the spike do not hinder the mechanism by which it binds to the enzymatic domain of ACE2^3^. This evolutionary space enables the virus to evolve quickly in response to both the natural immune response and biomedical interventions, rendering therapeutic antibodies and vaccines less effective across variants[2, 3, 7]. Additionally, given the large number of prophylactics and treatments that focus on this mechanism of entry for the virus, significant selection pressure is placed on the spike protein, increasing the risk of evolutionary escape by the virus from treatment [2, 3, 7]. It is thus imperative to search for antiviral treatments that target novel mechanisms in viral cell entry, to reduce the rate of transmission. As such treatments would provide an orthogonal selection pressure on the virus, they would provide an additional evolutionary constraint for the virus, thereby slowing down its rate of evolution, which would increase the durability of both natural and vaccinal immunity,

The 78 kilo-Dalton glucose-regulated protein (GRP78), also known as binding immunoglobulin protein (BiP), is involved in the stress response of the endoplasmic reticulum[9, 10]. GRP78 has also been shown to be a host-auxiliary factor for SARS-CoV-2 entry and infection, and therefore offers promise as a target for novel antiviral treatments[11]. Specifically, A.J Lee and coworkers showed that the substrate binding domain (SBD) of GRP78 can form a complex with the SARS-CoV-2 spike protein and ACE2 in cells, and that *in vitro* GRP78 directly binds to the receptor binding domain (RBD) of SARS-CoV-2 spike protein and ACE2. GRP78 has two binding domains, the nucleotide binding domain (NBD) and the SBD, with multiple X-ray structures available in the Protein Data Bank (PDB)[12]. Previous protein docking studies have suggested that the *β*region of the SBD, the spike protein RBD region IV (residues 480 – 488), and ACE2 form a ternary complex [10, 13]. It is anticipated that a small molecule that disrupts the binding of the GRP78 SBD to the spike protein RBD, will prevent the formation of the GRP78- spike RBD-ACE2 complex, thereby preventing viral entry into the cell. The identification of small molecules that bind to the SBD domain of GRP78 and disrupt its binding to the spike RBD could, therefore, pave the way for the development of antivirals for early treatment of SARS-CoV-2[10] and potentially other coronaviruses. Disrupting GRP78 function in the host can be expected to have toxicities associated with it. Such potential toxicities could be alleviated by developing a GRP78 inhibitor for intranasal delivery—the local administration of small quantities of drug to the nasal mucosa is likely to limit its side-effect potential.

*In vitro* studies have also demonstrated that disruption of GRP78 function downregulates ACE2 expression[13]. This effect may be due to the well-characterized chaperone activity of GRP78, and potentially could be mediated by the binding of a small molecule in the active site of GRP78 in its NBD. Small molecules with dual binding to both the GRP78 SBD and the NBD sites may therefore be even more effective.

Antivirals that target GRP78 instead of the spike RBD directly would place orthogonal selection pressure on the virus and limit the development of resistance to therapeutic antibodies and vaccines. Additionally, as GRP78 is a host protein, disrupting its interaction with ACE2 makes it more challenging for SARS-CoV-2 to evolve resistance to the GRP78 inhibitor itself. If deployed widely, a GRP78-targeting antiviral could significantly slow the spread of the virus by directly preventing infection (if used as an early-exposure prophylactic or early treatment option) and by increasing the duration of effectiveness of both the natural and vaccinal immune response.

In this work, we set out to develop an accurate model of the GRP78-spike RBD-ACE2 complex through computational protein-protein docking methods, and to use this model together with computationally mapped hot spots on the surface of the GRP78 SBD to identify the RBD binding pocket on the SBD of GRP78. This putative binding site was then used to screen for novel small molecule binders expected to bind to GRP78 and prevent its binding to the spike RBD. We anticipated that numerous molecules with properties that make them suitable for oral and/or intranasal administration could be computationally demonstrated to bind to the spike RBD binding site of GRP78 and prevent the formation of the GRP78-spike RBD-ACE2 complex. The results of this work could serve as a starting point for the further development of effective treatments against SARS-CoV-2. Additionally, the information gained about the druggability of the SBD binding pocket of GRP78 could also be used as a starting point for the development of antivirals against a broad spectrum of coronaviruses that utilize a similar mechanism for viral cell entry.

## Methods

### GRP78-Spike RBD-ACE2 Complex Model Generation

#### Protein-protein docking

A model of the GRP78-spike RBD-ACE2 complex was generated utilizing the HADDOCK Web Server[14, 15] version 2.4. We docked the GRP78 structure to the spike protein RBD-ACE2 complex structure. Structures of the ATP-bound state of human GRP78 (PDB ID: 5E84) and of the SARS-CoV-2 spike protein RBD-human ACE2- B^0^AT1 complex (PDB ID: 6M17) were utilized. For GRP78, chain A of the 5E84 structure with all waters, ions, and ATP removed, and was designated as Molecule 1 in the Haddock submission. For the spike protein RBD-ACE2 complex, chain B (ACE2) and E (spike protein RBD) with all waters and ions removed was designated as Molecule 2. Residues in chain E were renumbered to start at 633 such that HADDOCK would recognize spike RBD-ACE2 as one “molecule” during the docking.

Models of the multimeric complex have been previously published[9, 10] but have not been made publicly available. We sought to reproduce the HADDOCK model of the complex[13] and to make it freely available as a resource to others interested in developing inhibitors for this complex. “Active residues” were specified for both Molecule 1 and 2 for the HADDOCK docking. The “active residues” input for the spike RBD-ACE2 complex (Molecule 2) were C480 to C488 of the spike RBD as had previously been done. Pep42 is a cyclic peptide (CTVALPGGYVRVC)[10, 16] previously determined to be a GRP78 ligand[17]. The spike RBD C480 to C488 region (CNGVEGFNC) is considered “cyclic” due to the disulfide bond formed between the two cysteine residues and is the most homologous region of the spike RBD to Pep42 with three identical residues (see Fig. 1). For GRP78 the active residues were the ones identified as those H-bonding to the spike RBD in the model generated by Ibrahim et al.[13], specifically GRP78: T428, V429, S452, and T458, all of which are located in the substrate binding site of GRP78 SBD. [12] All other HADDOCK parameters were left at the default values. The models from the top scoring clusters, those with lowest HADDOCK scores, were then visualized in Maestro Version 12.9.137 (Schrödinger Release 2022-2) (Schrödinger, LLC, New York, NY, 2021) and compared to the one presented by Ibrahim et al.[10].

**Fig. 1.**
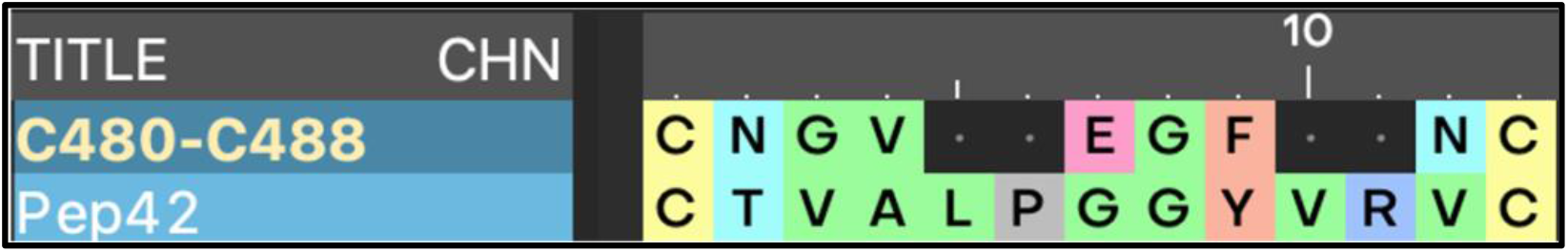
Sequence Alignment of C480-C488 Spike RBD Region and Pep42

#### FTMap GRP78 hot spot identification

As an orthogonal approach for identifying the spike RBD binding site on GRP78, hot spots were computationally mapped on the surface of the GRP78 SBD using the FTMap[18] server. FTMap is a computational mapping server that identifies regions of the structure surface that are expected to make a major contribution to the ligand binding free energy. [19] FTMap docks probes and scores the probe poses using detailed energy expression, identifying regions which bind clusters of multiple probe types as binding hot spots.[19] The structure of GRP78 (5E84 chain A) with the NBD (residues 1 to 400) deleted was used. All other parameters were left as the defaults. Hot spots corresponding “druggable” binding sites, those containing 16 or more probe clusters, were visualized [20].

#### Binding pocket and residues identification

The putative spike binding site on GRP78 was also visualized utilizing the Creating Binding Site Surfaces tool in Maestro, which displays a surface on the receptor and the ligand in the interaction region. The active residues on GRP78, the spike RBD, and ACE2 were established using the Protein-Protein Interaction tool, which identifies all residues located within 5 Å of the selected protein in Maestro as well.

### Structure-Based Virtual Screening

#### Protein and grid preparation

The structure of GRP78 (5E84 chain A) was prepared using the Protein Preparation Workflow Tool in Maestro with default parameters. Glide version 12.9.137 (Schrödinger, LLC, New York, NY, 2021)[21–23] molecular docking grids for GRP78 were generated by specifying the centroid as the residues of GRP78 within 5Å of the Spike RBD C480- C488 region in our GRP78-RBD-ACE2 complex model (GRP78: I426,T428, V429, V432, T434, K435, G449, I450, P451, S452, V453). Multiple receptor grids were generated by iteratively increasing the size of the grid until acceptable results were obtained for the control docking experiment described below. The final size of the grid was 11 × 13 × 23 Å for the ligand diameter midpoint box and 18 × 18 × 18 Å for the docked ligand maximum length box. Otherwise, default parameters were utilized.

#### Control Molecule Docking

To ensure that the receptor grid was adequate for conducting the virtual screens and to have a threshold value to compare our docking scores against we first docked the “cyclic” C480-C488 region of the spike RBD as a control. The structure of the C480- C488 peptide was taken from the spike protein RBD-ACE2 complex structure (PDB ID: 6M17), the peptide termini were manually capped with N-methyl (NME) and acetyl (ACE) residues, respectively, and the peptide was prepared using the Protein Preparation Workflow tool in Maestro. Default settings were used for all parameters. The peptide was docked utilizing Glide with SP scoring [21–23], and the resulting poses were compared to the position of the RBD C480- C488 region in the GRP78-Spike RBD-ACE2 complex model described above.

#### Preparation of ligand libraries for docking

Three ligand libraries from ZINC15[24] were prepared for docking: Microsource World Drug Collection (MWDC), NIH Clinical Collection (NCC), and ChEMBL Database. MWDC is an extension of the US Drug Collection database, which includes 400 drugs that have been marketed or are currently marketed in Europe and Asia. NCC consists of those compounds that previously had advanced to clinical trials[25]. Lastly, ChEMBL consists of bioactive drug-like small molecules, and the database includes calculated chemical properties and abstracted bioactivities[26]. For MWDC and NCC, the libraries were downloaded in SDF format and prepared for docking using the LigPrep tool in Maestro with default parameters. For ChEMBL the dataset was filtered prior to downloading by “for sale” and “clean”[24], and then similarly processed with LigPrep. Prior to docking, all sets were further filtered based on physicochemical and ADME properties calculated using QikProp[27]. The properties and thresholds used are given in Table 1 and were chosen to select compounds expected to be orally bioavailable.

**Table 1.** Filtering Criteria for Ligand Libraries

| Name | Abbreviation In Maestro | Range/Value |
| --- | --- | --- |
| ***(i) Oral Drug Properties*** |  |  |
| *Octanol/Water Partition Coefficient* | *QPlog Po/w* | - 2 to 6.5 |
| *Aqueous Solubility* | *QPlogS* | -6.5 to 0.5 |
| *Caco-2 Cell Permeability* | *QPPCaco* | > 25 |
| *MDCK Cell Permeability* | *QPPMDCK* | > 25 |
| *Percent Human Oral Absorption Values* | *HumanOralAbsorb* | > 50% |
| ***(ii) Additional Filters for ChEMBL Library Screening*** |  |  |
| *Number of Reactive Functional Groups* | *#rtvFG* | 0 |
| *Molecular Weight* | *mol_MW* | < 600 |
| *Number of Violations of Jorgensen’s Rule of Three<sup>a</sup>* | *RuleOfThree* | 0 |

<sup>a</sup> Rule of Three: *QPlogS* > -5.7, *QPPCaco* > 22 nm/s, # Primary Metabolites < 7.[28]

#### Docking ligands and analysis of poses

Glide with SP scoring [21–23] was used to dock the ligand libraries with default settings. Docking poses (up to 5 per molecule) were visually inspected in Maestro. Those that overlapped with the position of the RBD C480-C488 region in the GRP78-RBD-ACE2 complex model and with top-ranking hot spot in substrate binding region of the SBD of GRP78 were selected for further analysis. For MWDC and NCC, all docked ligands were examined, while for ChEMBL only the top 2000 based on the Glide docking score were. For each selected compound, the pose with the lowest Emodel score[21, 22] was retained for subsequent clustering.

#### Clustering and identification of known actives

The selected docked molecules from all three libraries were combined and clustered utilizing the Fingerprint Similarity and Clustering tool within Canvas[29, 30]. The Fingerprint settings chosen were the following: 32-bit precision, linear fingerprint type, daylight invariant atom types and bonds distinguished by bond order atom typing schemes. The similarity metric specified was Tanimoto and the centroid linkage method was chosen for clustering. The SARS-CoV-2 Screening Data 2020-21 in the ChEMBL Database was searched for all clustered compounds and active compounds with half maximal inhibitory concentration (IC50) <= 10 *μ*M or between 10 *μ*M and 50 *μ*M were identified.

## Results

### GRP78-Spike RBD-ACE2 Complex Model

As described in the methods section, a model of GRP78 bound to the SARS-CoV-2 spike RBD-ACE2 complex was generated. Models in the top ranked cluster output by Haddock were visualized and the best scoring model was selected (Fig. 2); the HADDOCK score was -77.9 +/- 3.7. Our model is similar to the one described by Ibrahim et al.[13] in terms of score (77.0 +/- 3.1) and overall topology but is distinct with respect to specific interactions. Our model identifies the interacting residues of GRP78 as I426, T428, V429, V432, T434, G449, I450, P451, S452, V453, predominantly situated in the *L*1,2 and *L*3,4 loops on the SBD*β*of GRP78, which surround the substrate binding site.[12] The interacting residues from the spike RBD were T478, A481, G482, V483, G484, G485, P486, A487, respectively. This is consistent with the cyclic region “C480 to C488” being the region of the spike RBD predicted, based on homology to the known GRP78 ligand, Pep42, to interact with GRP78 [17]. Consequently, the binding pocket for spike RBD was located on the SBD of GRP78 as expected (*Fig. 3*).

**Fig. 2.**
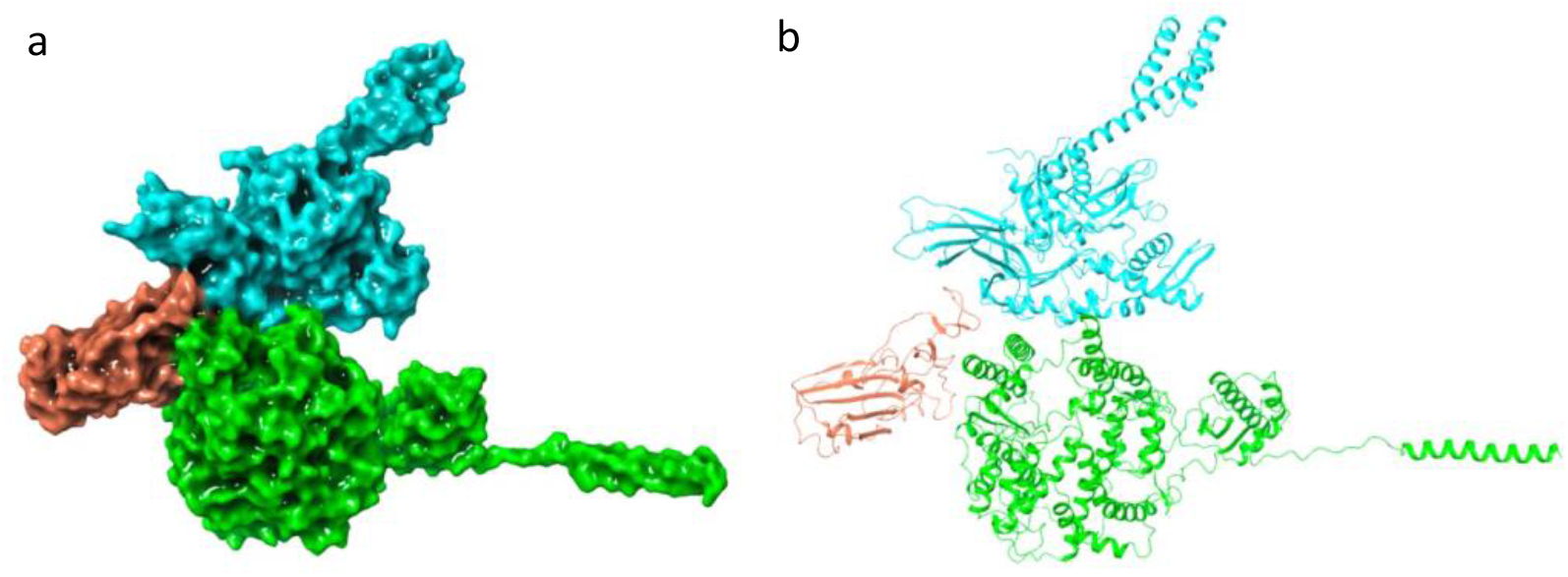
Model of the GRP78-Spike RBD-ACE2 Complex. GRP78 is shown in light blue, Spike RBD in orange, ACE2 in green, in (A) in a surface representation and in (B) a in cartoon representation

**Fig. 3.**
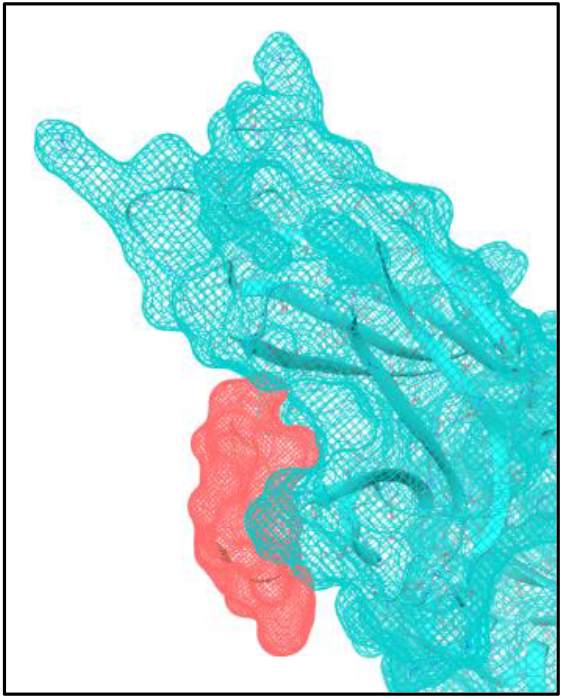
C480 - C488 region of spike RBD bound to a pocket of on the surface of the SBD of GRP78 in the GRP78-spike RBD-ACE2 model. C480-C488 of the spike RBD is represented by a molecular meshgrid (generated using the Creating Binding Site Surfaces tool in Maestro) in corale while GRP78 is in cyan

### FTMap mapping of GRP78 SBD domain

Hot-spot mapping of the GRP78 SBD was performed to confirm independently the location of the spike RBD binding site. Three hot spots consisting of 16 or more probe clusters were identified (see Fig. 4). The top ranked hot spot (with 22 probe clusters) found on the SBD surface is located at the NBD-SBD domain interface, essentially identifying the NBD binding site on the SBD surface. Second top ranked hot spot (with 18 probe clusters) is located far (> 18 Å) from the substrate binding site where the spike RBD is expected to bind. The third ranked hot spot (with 17 probe clusters) is located in the substrate binding site. According to Kozakov et al. [20], the likelihood that a binding site hot spot is “druggable” increases if it consists of 16 or more probe clusters, and if there is one or more secondary hot spots within 8 Å of it, creating a cluster ensemble with a maximum dimension of at least 10 Å. The third ranked hot spot (shown in yellow in Fig. 4) has a center-to-center distance of less than 8 Å to two other hot spots (represented in red and blue, respectively) and the maximum dimension of the cluster ensemble is slightly over 10 Å. Furthermore, this hot spot is consistent with the position of the spike RBD binding site in our HADDOCK model.

**Fig. 4.**
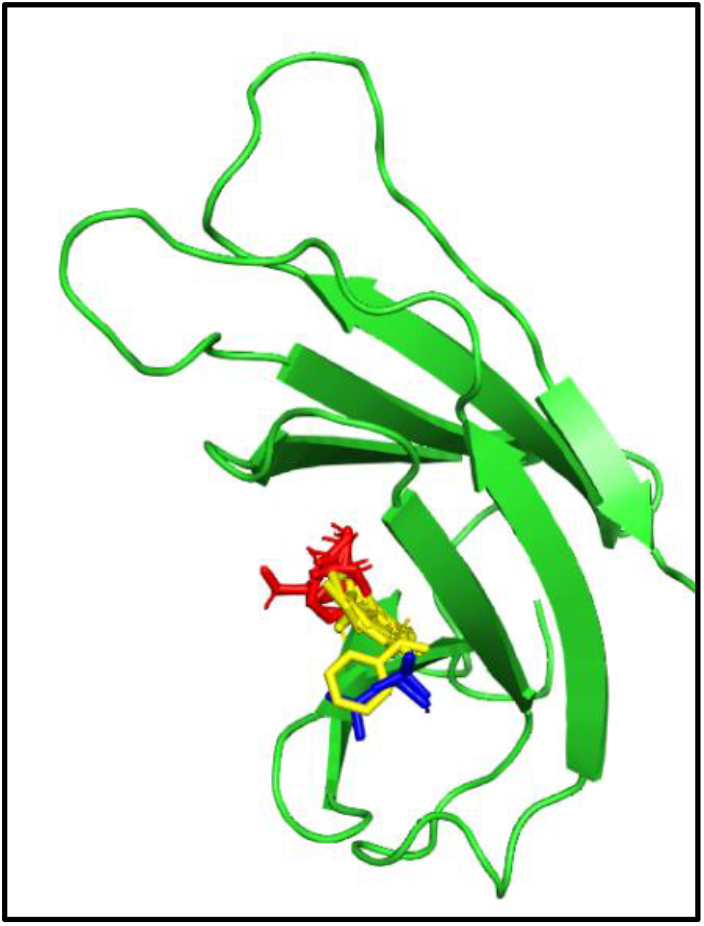
GRP78 SBD represented in green cartoon shown with probes in the third ranked hot spot shown in ball-and-stick in yellow. Adjacent hot spots are shown in red (the 4^th^ ranked) and blue (the 8^th^ ranked)

### Structure-Based Virtual Screening

As described in the methods, the capped C480 - C488 control peptide was first docked to GRP78 as a control; 28 poses were generated by Glide with docking score less than 0 kcal/mol. Pose number 18 was most similar to the position of C480-C488 in the spike RBD in the GRP78- spike RBD-ACE2 complex model had a docking score of -2.275 kcal/mol. Using the same setup as for the control peptide, the WDC, NCC, and ChEMBL compound libraries (filtered as described in the Methods section) were screened for binders to the putative spike RBD binding pocket on the SBD of GRP78.

Docking poses were visually inspected and only those that overlapped with both the C480- C488 region of the spike RBD in the model of the complex, as well as with probes in the corresponding FTMap hot spot, were retained (see an example in Fig. 5). The screen resulted in 23 hits with docking scores ranging from -5.50 to -0.43 kcal/mol from the WDC library, 42 with docking scores ranging from -4.70 to -1.41 kcal/mol from the NCC library, and 79 hits with docking scores ranging from -6.28 to -5.45 kcal/mol from the 2000 top-scoring compounds from the ChEMBL library (see Table 2).

**Table 2.** Summary of Docking Results

| Database Name | Hits | Molecules Downloaded | Conformers Docked | Poses Generated | Maximum Docking Score (kcal/mol) | Minimum Docking Score (kcal/mol) |
| --- | --- | --- | --- | --- | --- | --- |
| MWDC | 23 | 301 | 442 | 435 | -4.947 | -0.431 |
| NCC | 42 | 671 | 1093 | 1086 | -4.704 | -0.304 |
| ChEMBL | 79 | 479,516 | 697,147 | 402,808 | -6.278 | -5.454 |

**Fig. 5.**
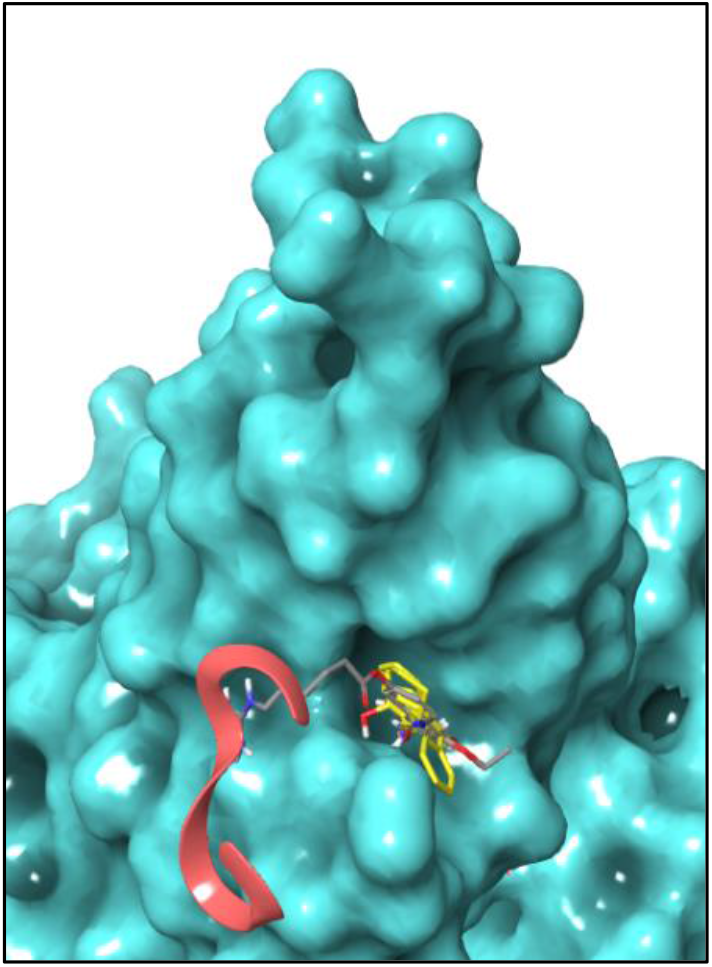
An example of a docking hit (ZINC000002002226) overlapping with the position of the C480-C488 region of the spike protein RBD in the model or the GRP78-RBD-ACE2 complex and probes from the calculated hot spot. The docking hit is shown in thick stick colored by element with carbons in grey, the C480 - C488 region is shown in a salmon ribbon, probes from the hot spots are in yellow, and the and GRP78 structure is shown in cyan in a surface representation

Fingerprint clustering of the 144 hits produced 12 clusters (see Table 3), and the 2D structure of cluster representatives are shown Fig. 6. Cluster 5 consists of 131 molecules and the other clusters each included no more than two molecules. Approximately 39% of these virtual screening hits across the clusters were present in the SARS-CoV-2 Screening Data 2020-21 set in ChEMBL, which is a repository of data combined from various small-molecule repurposing screens to identify inhibitors of SARS-CoV-2 viral activity. None of the hits were found among the compounds in the set with IC50s between 10 *μ*M and 50 *μ*M. Notably, however, seven compounds from Cluster 5 inhibited SARS-CoV-2 with IC50 < 10 *μ*M based the SARS-CoV-2 Screening Data 2020-21 (Table 4).

**Table 3.** Results of Fingerprint Clustering Performed on Docking Hits

| Cluster Number | Cluster Size | ChEMBL Molecules | NIH Molecules | World Molecules | Centroid Representative | Representative From | MW |
| --- | --- | --- | --- | --- | --- | --- | --- |
| 1 | 1 | 1 | 0 | 0 | ZINC000018173509 | ChEMBL | 395.4 |
| 2 | 1 | 1 | 0 | 0 | ZINC000013130700 | ChEMBL | 324.3 |
| 3 | 1 | 0 | 1 | 0 | ZINC000003881556 | NIH Clinical Collection | 356.5 |
| 4 | 1 | 0 | 0 | 1 | ZINC000000607861 | World Drug Collection | 377.5 |
| 5 | 131 | 76 | 37 | 18 | ZINC000002861721 | ChEMBL | 409.3 |
| 6 | 2 | 0 | 1 | 1 | ZINC000001542890 | NIH Clinical Collection | 338.4 |
| 7 | 1 | 0 | 1 | 0 | ZINC000004245630 | NIH Clinical Collection | 406.5 |
| 8 | 1 | 0 | 1 | 0 | ZINC000003921872 | NIH Clinical Collection | 412.6 |
| 9 | 1 | 1 | 0 | 0 | ZINC000006521422 | ChEMBL | 274.3 |
| 10 | 1 | 0 | 0 | 1 | ZINC000000603195 | World Drug Collection | 360.4 |
| 11 | 1 | 0 | 1 | 0 | ZINC000003920673 | NIH Clinical Collection | 467.0 |
| 12 | 2 | 0 | 0 | 2 | ZINC000003861133 | World Drug Collection | 394.5 |

**Table 4.**
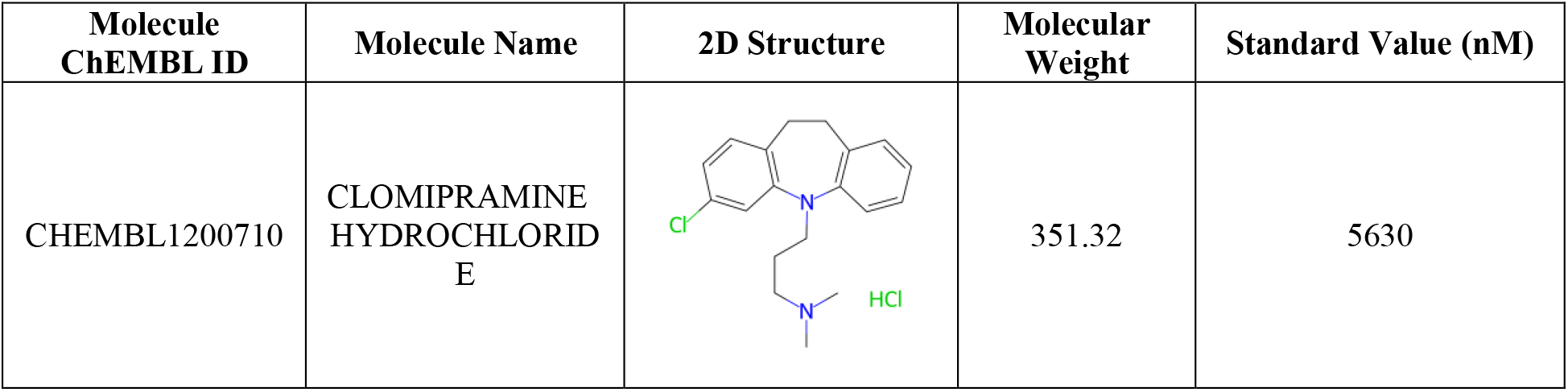
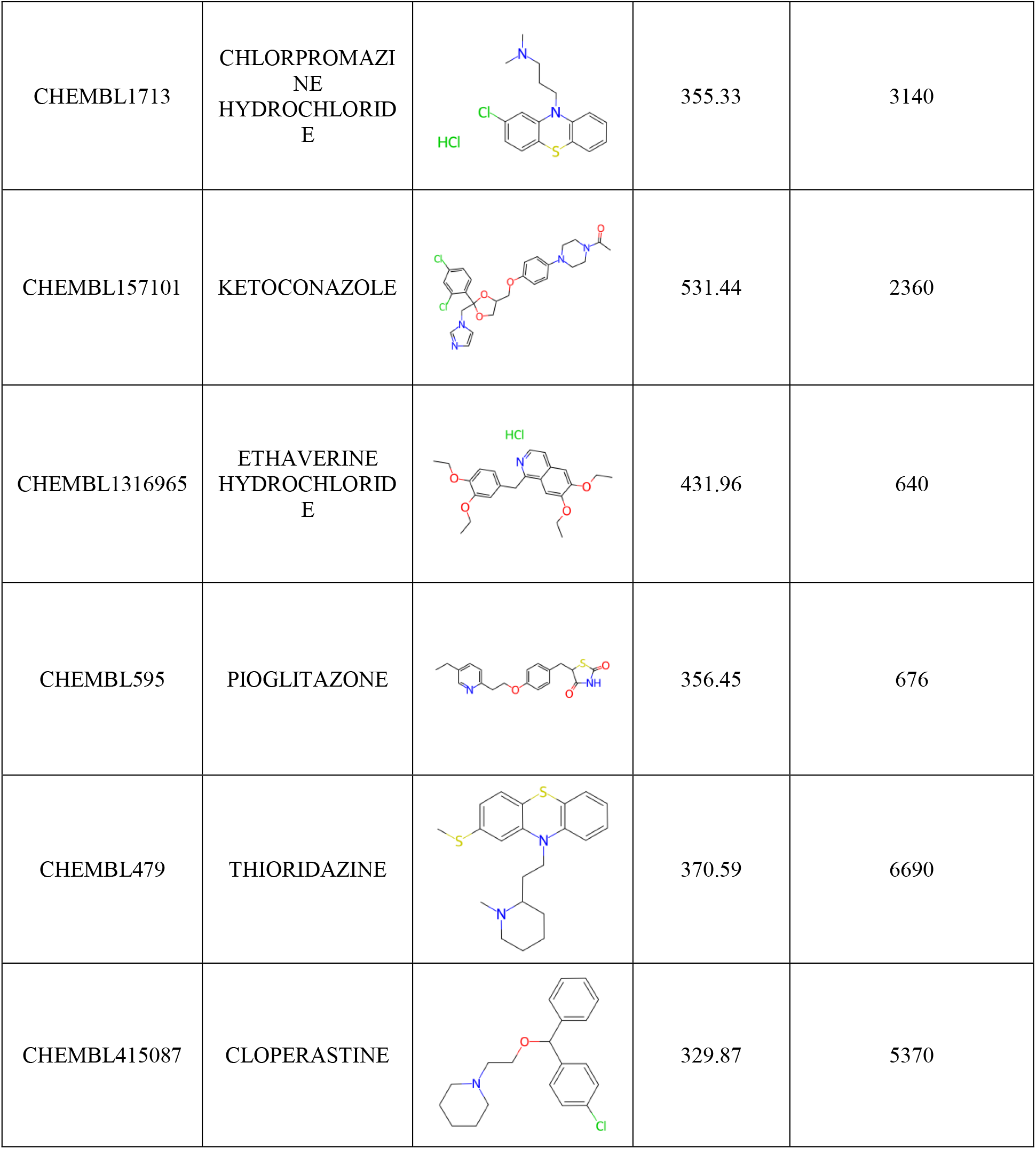
Docking Hits with an IC50 below 10 *μ*M in the ChEMBL SARS-CoV-2 Screening Data 2020-2021

| Molecule ChEMBL ID | Molecule Name | 2D Structure | Molecular Weight | Standard Value (nM) |
| --- | --- | --- | --- | --- |
| CHEMBL1200710 | CLOMIPRAMINE HYDROCHLORIDE |  | 351.32 | 5630 |
| CHEMBL1713 | CHLORPROMAZINE<br>HYDROCHLORIDE |  | 355.33 | 3140 |
| CHEMBL157101 | KETOCONAZOLE |  | 531.44 | 2360 |
| CHEMBL1316965 | ETHAVERINE<br>HYDROCHLORIDE |  | 431.96 | 640 |
| CHEMBL595 | PIOGLITAZONE |  | 356.45 | 676 |
| CHEMBL479 | THIORIDAZINE |  | 370.59 | 6690 |
| CHEMBL415087 | CLOPERASTINE |  | 329.87 | 5370 |

**Fig. 6.**
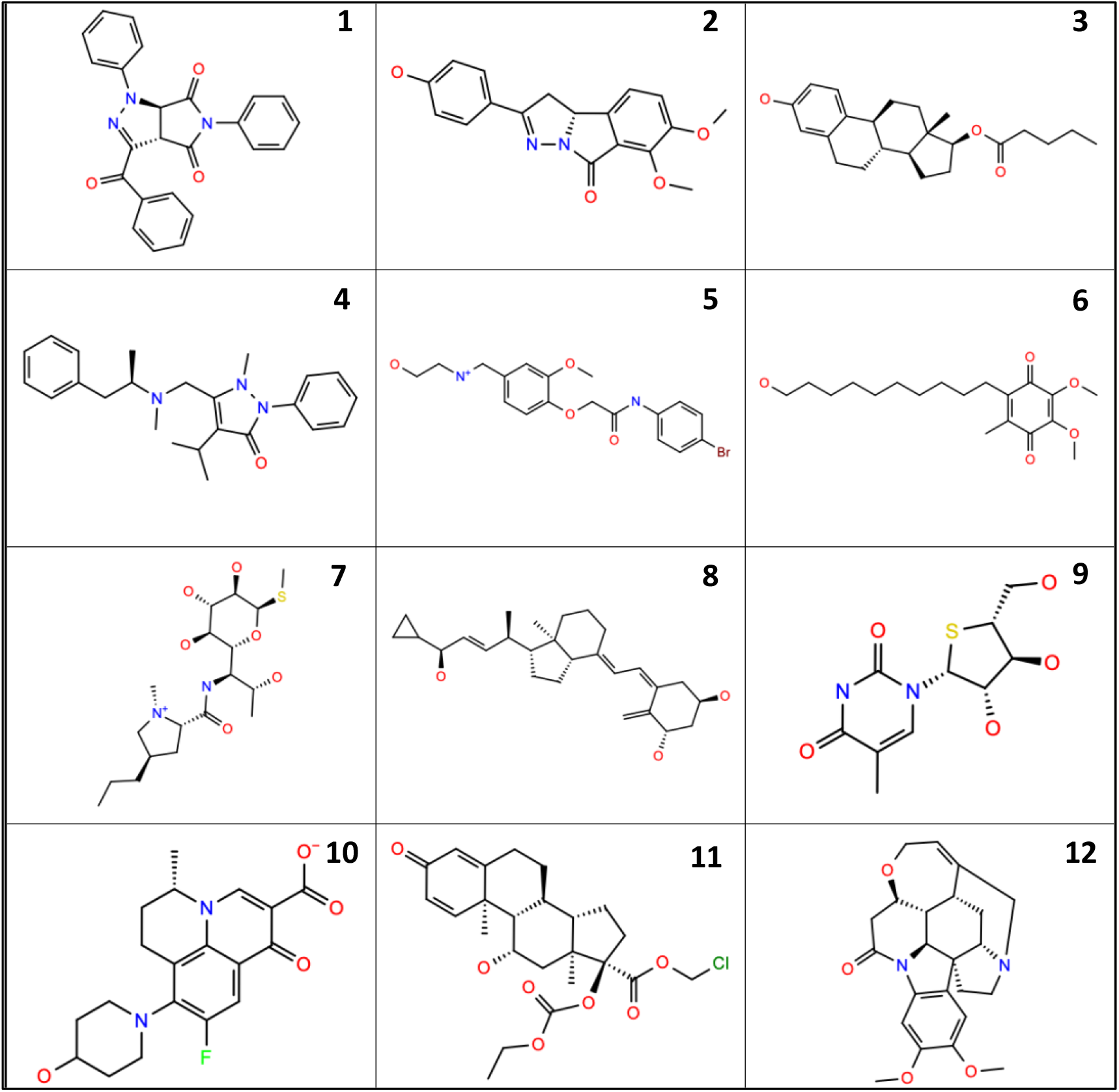
2D structures of cluster centroids from each cluster (as indicated by the number)

## Discussion

GRP78 has been shown to act as a secondary receptor responsible for host cell entry with GRP78 binding to the SARS-Cov-2 Spike protein and human ACE2 [11]. Our model of the GRP78-Spike RBD-ACE2 complex created in HADDOCK is consistent with previously generated models[13], and shows that the main interaction between the Spike RBD and GRP78 occurs between the C480-C488 region and substrate binding site of GRP78. We identified the active residues involved in the interaction between the Spike protein and GRP78, observing some differences compared to previously published models. Furthermore, hot spot mapping of the GRP78 SBD surface, identified a binding site at the location on GRP78 where the Spike RBD binds in our model of the complex that is predicted with high confidence to be “druggable”. In addition, the control “cyclic” peptide representing the “C480-C488” region of the spike RBD docked to GRP78 at that binding site with a favorable docking score. These results taken together strongly point to this binding site as the putative location where the Spike RBD binds to GRP78 when in complex with ACE2. The results also indicate that this binding site on GRP78 is a druggable site, is suitable for virtual screening, and that small molecules binding to GRP78 at this site should disrupt complex formation.

From the virtual screens, 144 hits with drug-like properties and favorable docking scores were identified that would be expected to prevent the formation of the GRP78-Spike RBD-ACE2 complex. The ChEMBL hits had more favorable docking scores than the hits from MWDC and NCC due to the much larger size of that database. If more than the 2000 top scoring molecules from the ChEMBL database had been visualized, additional hits with slightly less favorable scores would almost certainly been identified.

After clustering, 12 clusters representing at least 12 chemotypes were identified. The 65 hits from MWDC and NCC are either drugs approved by some countries or molecules that reached clinical trials for other purposes. Cluster 5 consisted of 76 hits from ChEMBL, 18 from WDC, and 37 from NCC. In addition, a maximum common substructure search in the set showed that there are at least four substructures of at least 20 compounds based on common structure (Supplementary Table 2).

Seven of these hits from ChEMBL are actives based on the SARS-CoV-2 Screening Data 2020-21 in ChEMBL. The mechanism of action of these hits against SARS-CoV-2 is unknown, suggesting that it is possible that the activity of these compounds could be due to GRP78 binding. Of the 7 SARS-CoV-2 active hits identified, 2 are phenothiazines (Chlorpromazine and Thioridazine) and one is a dibenzazepine tricyclic antidepressant (Chlomipramine), which is structurally and chemically related to phenothiazines[31].

Since GRP78 is a human target that plays an important role in cellular function[32], systematic disruption of its functions would not be desirable and could lead to unknown toxicities. The candidate molecules identified in the work, though, would not be expected to bind to the ATP binding site in the NBD of GRP78 and therefore are not likely to inhibit the ATPase activity of GRP78[33]. Blocking the substrate binding site of GRP78, as these hits are predicted to do, could possibly affect the chaperone activity of GRP78. Intranasal delivery of molecules capable of binding in the substrate binding site of GRP78, however, would be expected to limit toxicities, with very minimal systemic exposure expected. Most currently approved intranasally delivered drugs have properties similar to the hits identified in this work. The molecules identified herein are expected to diminish viral host cell entry via disruption of spike protein RBD-GRP78 interface and could pave the way for the future development of novel antiviral treatment options for SARS-CoV-2 and potentially other viral pathogens.

## Supporting information

GRP78-spike RBD-Ace2 complex model

## Acknowledgments

We thank Kenji C. Walker for thoughtful discussions related to the properties of intranasally delivered drugs.

