## Supplemental Data for "Identification of a Druggable Site on GRP78 at the GRP78-SARS-CoV-2 Interface and Compounds to Disrupt that Interface"

**Supporting Information: Identification of a Druggable Site on GRP78 at the Putative GRP78-SARS-CoV-2 Interface and Virtual Screening of Compounds to Disrupt that Interface**

Maria Lazou[1], Jonathan Hutton[1], Arijit Chakravarty[2], Diane Joseph-McCarthy[1]*

[1]Department of Biomedical Engineering, Boston University

[2]Fractal Therapeutics

*Table 1. Oral Results for Virtual Screening*

| ZINC ID | Database | Canvas Cluster Index | Docking Score |
|---|---|---|---|
| ZINC000001574876 | ChEMBL | 5 | -6.278 |
| ZINC000040395734 | ChEMBL | 5 | -6.124 |
| ZINC000003222846 | ChEMBL | 5 | -6.072 |
| ZINC000040395471 | ChEMBL | 5 | -6.03 |
| ZINC000003266556 | ChEMBL | 5 | -6.03 |
| ZINC000000279195 | ChEMBL | 5 | -6.007 |
| ZINC000000003962 | ChEMBL | 5 | -6.005 |
| ZINC000004394380 | ChEMBL | 5 | -5.989 |
| ZINC000001996784 | ChEMBL | 5 | -5.98 |
| ZINC000008609343 | ChEMBL | 5 | -5.946 |
| ZINC000004825401 | ChEMBL | 5 | -5.942 |
| ZINC000006521422 | ChEMBL | 9 | -5.924 |
| ZINC000005095303 | ChEMBL | 5 | -5.882 |
| ZINC000000493815 | ChEMBL | 5 | -5.878 |
| ZINC000013601099 | ChEMBL | 5 | -5.876 |
| ZINC000000794244 | ChEMBL | 5 | -5.872 |
| ZINC000018229990 | ChEMBL | 5 | -5.863 |
| ZINC000013685662 | ChEMBL | 5 | -5.826 |
| ZINC000003189962 | ChEMBL | 5 | -5.819 |
| ZINC000002436579 | ChEMBL | 5 | -5.784 |
| ZINC000004644048 | ChEMBL | 5 | -5.76 |
| ZINC000009519770 | ChEMBL | 5 | -5.76 |
| ZINC000028950583 | ChEMBL | 5 | -5.748 |
| ZINC000015016427 | ChEMBL | 5 | -5.73 |
| ZINC000040639515 | ChEMBL | 5 | -5.721 |
| ZINC000001159058 | ChEMBL | 5 | -5.697 |
| ZINC000049785416 | ChEMBL | 5 | -5.697 |
| ZINC000009394479 | ChEMBL | 5 | -5.69 |
| ZINC000001427729 | ChEMBL | 5 | -5.673 |
| ZINC000004249247 | ChEMBL | 5 | -5.672 |
| ZINC000031870777 | ChEMBL | 5 | -5.659 |
| ZINC000013583589 | ChEMBL | 5 | -5.657 |
| ZINC000013378188 | ChEMBL | 5 | -5.653 |
| ZINC000002861721 | ChEMBL | 5 | -5.647 |
| ZINC000000204389 | ChEMBL | 5 | -5.647 |

| | | | |
|---|---|---|---|
| ZINC000003960369 | ChEMBL | 5 | -5.64 |
| ZINC000018182321 | ChEMBL | 5 | -5.627 |
| ZINC000008855234 | ChEMBL | 5 | -5.607 |
| ZINC000014960224 | ChEMBL | 5 | -5.603 |
| ZINC000005084182 | ChEMBL | 5 | -5.603 |
| ZINC000005462119 | ChEMBL | 5 | -5.602 |
| ZINC000012471238 | ChEMBL | 5 | -5.598 |
| ZINC000012537137 | ChEMBL | 5 | -5.598 |
| ZINC000018173509 | ChEMBL | 1 | -5.597 |
| ZINC000001820502 | ChEMBL | 5 | -5.594 |
| ZINC000004249339 | ChEMBL | 5 | -5.594 |
| ZINC000012763896 | ChEMBL | 5 | -5.592 |
| ZINC000013472143 | ChEMBL | 5 | -5.573 |
| ZINC000013545030 | ChEMBL | 5 | -5.572 |
| ZINC000008670732 | ChEMBL | 5 | -5.564 |
| ZINC000013124992 | ChEMBL | 5 | -5.564 |
| ZINC000004479699 | ChEMBL | 5 | -5.563 |
| ZINC000002425514 | ChEMBL | 5 | -5.555 |
| ZINC000004755381 | ChEMBL | 5 | -5.554 |
| ZINC000001016079 | ChEMBL | 5 | -5.551 |
| ZINC000015055962 | ChEMBL | 5 | -5.546 |
| ZINC000006648255 | ChEMBL | 5 | -5.543 |
| ZINC000005620809 | ChEMBL | 5 | -5.54 |
| ZINC000028641215 | ChEMBL | 5 | -5.537 |
| ZINC000005580883 | ChEMBL | 5 | -5.533 |
| ZINC000000006846 | ChEMBL | 5 | -5.531 |
| ZINC000008619667 | ChEMBL | 5 | -5.516 |
| ZINC000009263941 | ChEMBL | 5 | -5.511 |
| ZINC000005940380 | ChEMBL | 5 | -5.5 |
| ZINC000008619635 | ChEMBL | 5 | -5.5 |
| ZINC000005060607 | ChEMBL | 5 | -5.492 |
| ZINC000013477102 | ChEMBL | 5 | -5.491 |
| ZINC000045353541 | ChEMBL | 5 | -5.49 |
| ZINC000005149064 | ChEMBL | 5 | -5.49 |
| ZINC000001248585 | ChEMBL | 5 | -5.489 |
| ZINC000064527339 | ChEMBL | 5 | -5.484 |
| ZINC000064560059 | ChEMBL | 5 | -5.481 |
| ZINC000000971866 | ChEMBL | 5 | -5.473 |
| ZINC000012074107 | ChEMBL | 5 | -5.467 |

| | | | |
|---|---|---|---|
| ZINC000002944477 | ChEMBL | 5 | -5.462 |
| ZINC000002229226 | ChEMBL | 5 | -5.458 |
| ZINC000022910125 | ChEMBL | 5 | -5.456 |
| ZINC000015941041 | ChEMBL | 5 | -5.456 |
| ZINC000013130700 | ChEMBL | 2 | -5.454 |
| ZINC000000001624 | MMWDC | 5 | -4.947 |
| ZINC000000603195 | MWDC | 10 | -4.713 |
| ZINC000000899213 | NCC | 5 | -4.704 |
| ZINC000000014669 | NCC | 5 | -4.509 |
| ZINC000003831004 | MWDC | 5 | -4.483 |
| ZINC000003921872 | NCC | 8 | -4.333 |
| ZINC000003831354 | MWDC | 5 | -4.259 |
| ZINC000001843099 | NCC | 5 | -4.201 |
| ZINC000000000654 | MWDC | 5 | -4.194 |
| ZINC000003875039 | MWDC | 5 | -4.024 |
| ZINC000000020248 | NCC | 5 | -4.004 |
| ZINC000003789437 | MWDC | 5 | -4.003 |
| ZINC000000968326 | NCC | 5 | -3.923 |
| ZINC000001530568 | NCC | 5 | -3.895 |
| ZINC000001542890 | NCC | 6 | -3.883 |
| ZINC000001542890 | MWDC | 6 | -3.883 |
| ZINC000000537795 | NCC | 5 | -3.867 |
| ZINC000003873160 | NCC | 5 | -3.856 |
| ZINC000001530695 | NCC | 5 | -3.789 |
| ZINC000000897251 | NCC | 5 | -3.78 |
| ZINC000003873781 | MWDC | 5 | -3.774 |
| ZINC000004245630 | NCC | 7 | -3.755 |
| ZINC000003871832 | NCC | 5 | -3.749 |
| ZINC000000044027 | NCC | 5 | -3.71 |
| ZINC000003799072 | NCC | 5 | -3.641 |
| ZINC000002015997 | MWDC | 5 | -3.623 |
| ZINC000000538065 | NCC | 5 | -3.599 |
| ZINC000003881556 | NCC | 3 | -3.551 |
| ZINC000000537805 | NCC | 5 | -3.462 |
| ZINC000000057252 | MWDC | 5 | -3.434 |
| ZINC000003995455 | NCC | 5 | -3.408 |
| ZINC000002015039 | MWDC | 5 | -3.372 |
| ZINC000003785268 | NCC | 5 | -3.369 |
| ZINC000001544805 | MWDC | 5 | -3.277 |

| | | | |
|---|---|---|---|
| ZINC000000968328 | NCC | 5 | -3.236 |
| ZINC000000607971 | NCC | 5 | -3.202 |
| ZINC000003830716 | NCC | 5 | -3.196 |
| ZINC000003920673 | NCC | 11 | -3.192 |
| ZINC000001530571 | NCC | 5 | -3.158 |
| ZINC000003786466 | NCC | 5 | -3.142 |
| ZINC000000057251 | MWDC | 5 | -3.093 |
| ZINC000003831578 | NCC | 5 | -3.086 |
| ZINC000003831578 | MWDC | 5 | -3.086 |
| ZINC000001542929 | NCC | 5 | -3.08 |
| ZINC000003775140 | NCC | 5 | -3.054 |
| ZINC000000643055 | NCC | 5 | -3.03 |
| ZINC000000968327 | NCC | 5 | -3.025 |
| ZINC000000608359 | MWDC | 5 | -3.01 |
| ZINC000001851149 | NCC | 5 | -2.936 |
| ZINC000001541570 | NCC | 5 | -2.837 |
| ZINC000001713761 | NCC | 5 | -2.665 |
| ZINC000001713761 | MWDC | 5 | -2.665 |
| ZINC000001530690 | NCC | 5 | -1.994 |
| ZINC000000607861 | MWDC | 4 | -1.945 |
| ZINC000001481844 | MWDC | 5 | -1.93 |
| ZINC000003873163 | NCC | 5 | -1.921 |
| ZINC000000643138 | NCC | 5 | -1.814 |
| ZINC000002002226 | NCC | 5 | -1.527 |
| ZINC000003792990 | NCC | 5 | -1.412 |
| ZINC000003872931 | NCC | 5 | -1.384 |
| ZINC000002015040 | MWDC | 5 | -1.229 |
| ZINC000000002181 | MWDC | 5 | -0.949 |
| ZINC000003861133 | MWDC | 12 | -0.629 |
| ZINC000003861134 | MWDC | 12 | -0.431 |
| ZINC000003812888 | NCC | 5 | -0.304 |

*Table 2. Results of Maximum Common Substructure Search on Cluster 5 Ligands. Icons created by SMARTSviewer [https://smarts.plus/]. Copyright: ZBH – Center for Bioinformatics Hamburg*

| SMARTS | 2D | Ligand Count |
|---|---|---|
| c1ccccc1CN(CCC2)c(n23)nc4c3c(=O)n(C)c(=O)n4C | 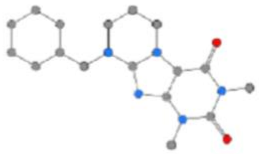 | 3 |
| c1ccccc1NC(=O)COc2ccc(C)cc2 | 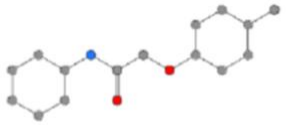 | 4 |
| c1ccccc1NC(=O)COc2ccccc2 | 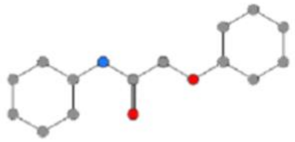 | 5 |
| cccccNCCOc1ccccc1 | 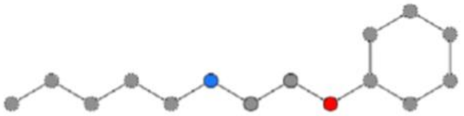 | 6 |
| c1ccccc1Cc2ccccc2 | 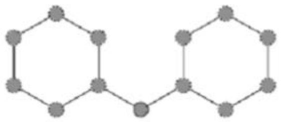 | 8 |
